# Developmental Mouse Brain Common Coordinate Framework

**DOI:** 10.1101/2023.09.14.557789

**Authors:** Fae A. Kronman, Josephine K. Liwang, Rebecca Betty, Daniel J. Vanselow, Yuan-Ting Wu, Nicholas J. Tustison, Ashwin Bhandiwad, Steffy B. Manjila, Jennifer A. Minteer, Donghui Shin, Choong Heon Lee, Rohan Patil, Jeffrey T. Duda, Luis Puelles, James C. Gee, Jiangyang Zhang, Lydia Ng, Yongsoo Kim

## Abstract

3D standard reference brains serve as key resources to understand the spatial organization of the brain and promote interoperability across different studies. However, unlike the adult mouse brain, the lack of standard 3D reference atlases for developing mouse brains has hindered advancement of our understanding of brain development. Here, we present a multimodal 3D developmental common coordinate framework (DevCCF) spanning mouse embryonic day (E) 11.5, E13.5, E15.5, E18.5, and postnatal day (P) 4, P14, and P56 with anatomical segmentations defined by a developmental ontology. At each age, the DevCCF features undistorted morphologically averaged atlas templates created from Magnetic Resonance Imaging and co-registered high-resolution templates from light sheet fluorescence microscopy. Expert-curated 3D anatomical segmentations at each age adhere to an updated prosomeric model and can be explored via an interactive 3D web-visualizer. As a use case, we employed the DevCCF to unveil the emergence of GABAergic neurons in embryonic brains. Moreover, we integrated the Allen CCFv3 into the P56 template with stereotaxic coordinates and mapped spatial transcriptome cell-type data with the developmental ontology. In summary, the DevCCF is an openly accessible resource that can be used for large-scale data integration to gain a comprehensive understanding of brain development.

## Introduction

Brain atlases provide a standard anatomical context to interpret brain structure^1,2^, function, neuronal connectivity^3–6^, molecular signature^7–9^, and cell type specific transcriptome data^10,11^. Atlases contain two related yet independent components: templates with distinct contrast features and anatomical delineations. Historically, mouse brain atlases existed in the form of 2D sections with histological staining and annotations based on cytoarchitecture often from a single male specimen^12,13^. These atlases present challenges in interpreting anatomical regions in 3D brains and may misrepresent anatomy across different individuals and sexes^14–16^. Moreover, recent advancements in cellular resolution whole mouse brain imaging techniques allow researchers to rapidly produce large scale 3D datasets^6,17–19^, calling for 3D reference atlases for data analysis^20,21^. Currently, the Allen Institute adult mouse brain common coordinate framework (CCFv3) serves as a 3D reference atlas to overcome many limitations of 2D atlases described above and to provide standardized spatial context to integrate data from different studies for the adult mouse brain^15^.

However, we lack high resolution 3D mouse brain atlases that account for developmental anatomy as well as ontologically consistent segmentations that can be used from fetal to adult brains. Developing brains undergo rapid shape and volume changes with cell proliferation and migration, often guided by regionally distinct gene expression^22–27^. Previously, the Allen Developing Mouse Brain Atlas (ADMBA) was created using a series of 2D histological sections with genoarchitecture guided segmentations based on the prosomeric model^24,28^. Although a recent study utilized computational methods to create 3D atlases from the 2D ADMBA^29^, it proves inadequate as the annotation methods used relied on label interpolation and edge detection of Nissl stained background data, which does not provide evidence for all developmental details. Moreover, existing MRI-based developmental mouse brain atlases do not meet community needs for cellular resolution 3D imaging and conflicting annotations across different atlases present major challenges to interpret brain areas across different studies^30–33^.

To overcome these limitations, we generated a developmental common coordinate framework (DevCCF) for mouse brains with expert-curated 3D segmentations at seven ages: embryonic day (E)11.5, E13.5, E15.5, E18.5, postnatal day (P)4, P14, and P56. Each age features undistorted morphology and intensity averaged symmetric templates from both male and female brains with at least four distinct magnetic resonance imaging (MRI) contrasts, as well as a light sheet fluorescence microscopy (LSFM) based template. Furthermore, we established developmentally consistent annotations based on the prosomeric model of mammalian brain anatomy^28^. We also integrated the CCFv3 and P56 DevCCF template, which enables users to compare the two distinct and complementary anatomical labels in the same space. We demonstrated the utility of the DevCCF by mapping GABAergic neurons and spatial transcriptome cell type data from developing and adult mouse brains, respectively. Lastly, we established an interactive viewer and downloadable datasets to freely distribute the DevCCF (https://kimlab.io/brain-map/DevCCF/).

## Results

### Pipeline overview to create the DevCCF

Creating each of the seven DevCCF atlases requires three primary steps: (1) 3D symmetric MRI and LSFM template generation, (2) multimodal template alignment (registration), and (3) anatomical segmentation. 3D symmetric templates for each modality are intensity and morphological averages of individual samples generated using non-linear registration with Advanced Normalization Tools (ANTs; Fig. 1a)^34^. We used LSFM to acquire high resolution images (up to 1.8 μm/voxel) with cell type specific labeling and other staining to acquire cellular features. We also used various MRI contrast datasets (e.g., diffusion weighted imaging; DWI) of *ex-vivo* in-skull mouse brain samples to create morphologically undistorted MRI templates with up to 31.5 μm isotropic voxel resolution (Fig. 1a). To mitigate morphological distortion due to removal of the brain from the skull and subsequent sample processing (e.g., tissue clearing), LSFM templates were registered to the MRI templates, generating multimodal DevCCF templates with cellular resolution features and undistorted morphology (Fig. 1a). Moreover, we established developmentally consistent anatomical segmentations based on cyto- and geno-architecture at seven developmental ages (E11.5, E13.5, E15.5, E18.5, P4, P14, and P56; Fig. 1b)^24,28,35^. Finally, we employed the DevCCF to map gene expression and other cell type information from various 2D and 3D imaging modalities to guide and validate our anatomical segmentations (Fig. 1c).

**Figure 1.**
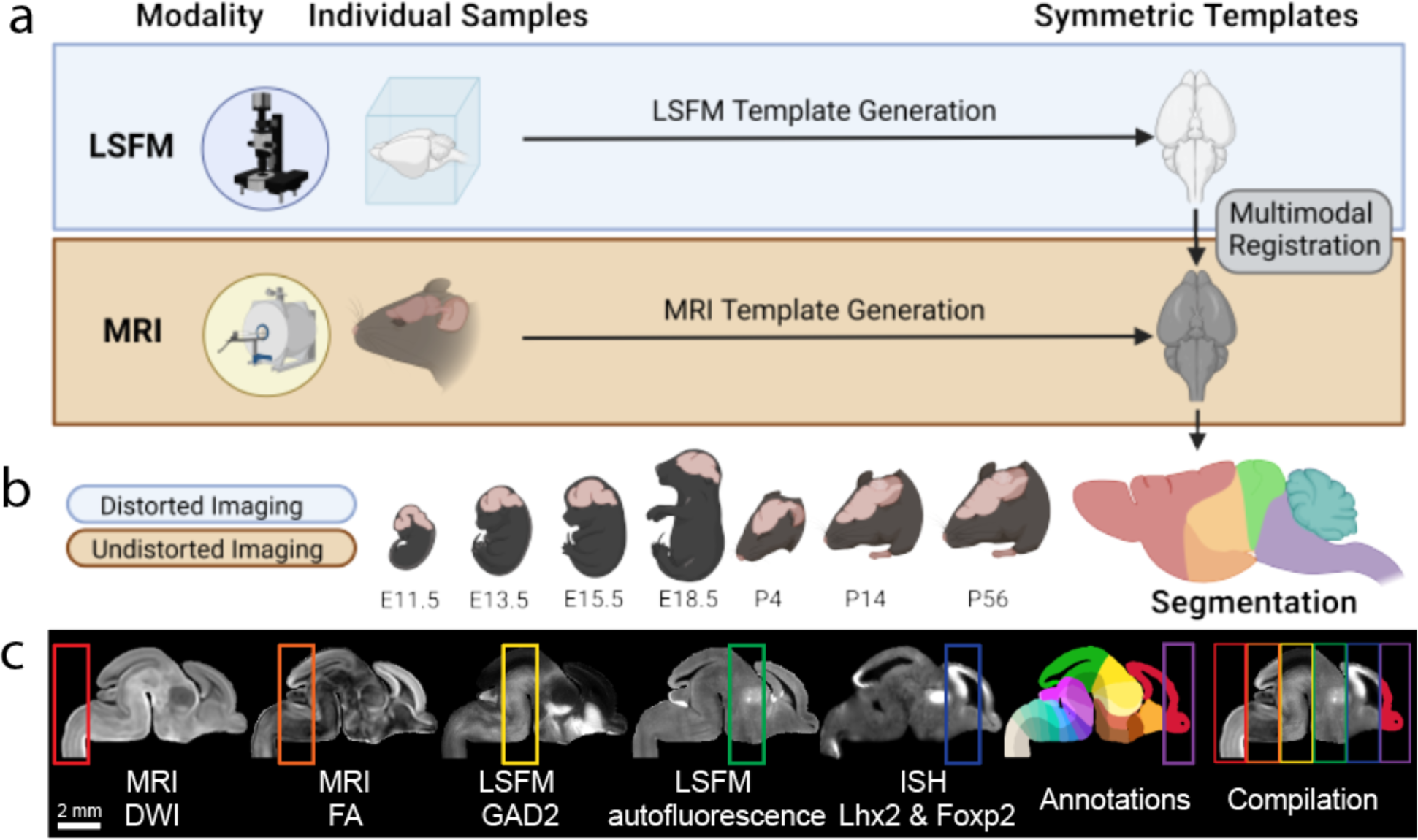
DevCCF overview. (a) Morphology and intensity averaged templates from LSFM (blue box) and MRI (yellow block) imaging. The LSFM template was aligned to the undistorted MRI template via multimodal registration. (b) Multimodal DevCCF with anatomical segmentations is established at four embryonic and three postnatal ages. (c) Sagittal slice of various E15.5 multimodal data registered to the E15.5 DevCCF, each highlighting unique anatomical features. From left to right: MRI DWI, MRI FA, LSFM GABAergic neurons from Gad2-Cre;Ai14 mice, LSFM autofluorescence, ISH data Lhx2 and Foxp2 gene expression to collectively guide DevCCF annotations.

### 3D Multimodal Developmental Mouse Brain Templates

We used MRI to image fixed *ex-vivo* in-skull samples at 31.5 μm (E11.5), 34 μm (E13.5), 37.5 μm (E15.5), 40 μm (E18.5), and 50 μm isotropic nominal voxel resolution (P4, P14, and P56) from both male and female mice (Fig. 2a). Using MRI DWI contrast for embryonic samples and apparent diffusion coefficient (ADC) contrast for postnatal samples, we iteratively registered age matched samples (n=6-14 samples per age; Extended Data Table 1) and their midline reflections to their composite average to create morphology and intensity averaged symmetric templates. We applied identical transformation fields and averaging procedures to each sample MRI contrast (transverse relaxation time weighted; T2-weighted, fractional anisotropy; FA, DWI, and ADC) to create a minimum of four symmetric MRI templates with distinct contrasts at each age (Fig. 2a). Moreover, we utilized tissue clearing and LSFM imaging to examine either whole heads (E11.5, E13.5, and E15.5) or brains (E18.5, P4, P14, and P56) at resolution of 1.8 (x) × 1.8 (y) × 5.0 (z) μm^3^/voxel. LSFM data was resampled to 10 μm isotropic voxel resolution and reflected across the midline to generate symmetrical templates as done in MRI templates (Fig. 2b).

**Figure 2.**
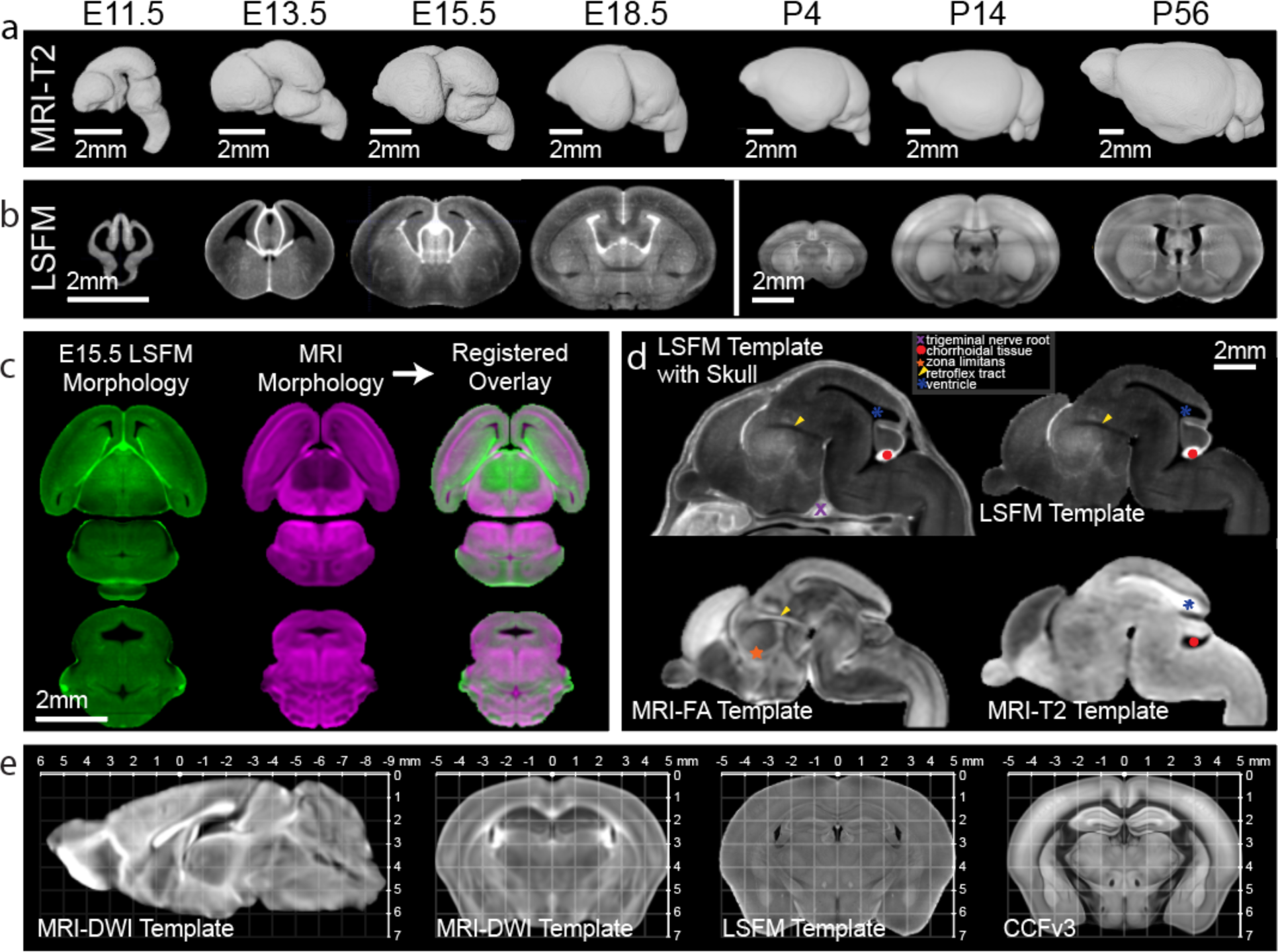
3D Multimodal Developmental Mouse Brain Templates. (a) 3D DevCCF MRI templates from T2-weighted contrasts. (b) DevCCF LSFM autofluorescence templates (coronal slice) before multimodal registration to the MRI template. (c) Multimodal registration of E15.5 LSFM to MRI DWI templates. Top: horizontal plane. Bottom: coronal plane. (d) DevCCF E15.5 multimodal templates with unique and complementary contrasts. Note distinct anatomical marks differentially highlighted. (e) The CCFv3 template is registered to the P56 DevCCF in stereotaxic coordinates. Left: MRI DWI template midline sagittal slice. Right: co-registered MRI DWI, LSFM autofluorescence, and CCFv3 coronal template coronal slices.

To establish the DevCCF with the benefits of both undistorted MRI morphology and the cellular resolution of LSFM in the same space, we optimized 3D landmark assisted multimodal registration methods to map LSFM templates to age matched MRI templates, enabling high resolution DevCCF templates across multiple modalities (Fig. 2c,d; see Methods for details). Templates with different contrasts or imaging modalities highlight distinct anatomic structures can be used independently or together for data viewing and registration (Fig. 2d). Moreover, we established the postnatal templates with stereotaxic coordinates by aligning bregma and lambda locations from MRI templates (Fig. 2e). This 3D coordinate information can facilitate use of the postnatal templates to design *in vivo* injections or recording experiments in the future^36,37^. Lastly, we conducted multimodal registration to establish the widely used CCFv3 template and its annotations in the stereotaxic system of the P56 templates to promote seamless integration of CCFv3 mapped data onto the new P56 DevCCF template and *vice versa* (Fig. 2e).

### 3D Anatomical labels based on a developmental ontology

Although distinct cytoarchitectures (e.g., with Nissl staining) have been useful to segment adult brains^12,13^, the same criteria are difficult to apply in developing brains due to immature and rapidly migrating cells. The 2D ADMBA is the current standard for the developing mouse brain atlas^24^. The ADMBA labels utilize an original developmental mouse brain ontology with up to 13-level hierarchical ontology based on the prosomeric model, which highlights genetically driven vertebrate brain developmental aspects^24,28^.

The DevCCF ontology provides an update to the ADMBA ontology to accommodate advancements in our understanding of developing vertebrate brain anatomy, such as concentric ring topology in the pallium (Extended Data Fig. 1)^35^. The DevCCF ontology also highlights data-driven modifications to the ADMBA ontology, such as renaming the isthmus to rhombomere 0^38^. Nevertheless, the DevCCF ontology closely follows the structure and information of the ADMBA ontology, matching region names, abbreviations, and color assignments (Supplementary Table 1)^24^.

To create anatomical segmentations, we outlined the surface of the brain using the T2-weighted MRI and LSFM templates, separating brain tissue from the skull and surrounding tissue in the templates. This included the brain surface that touches the skull, as well as folds in surface tissue where it touches itself. Landmark and theory guided segmentations continued level by level in line with the DevCCF ontology through at least level 5 for all ages (Fig. 3a). Segmentations adhere to prosomeric principles such as the understanding that each neuromere contacts only one neuromere caudally and one neuromere rostrally at any given location^28^. Finer segmentations continue past level 5 in most areas for all ages (Fig. 3b). We imported the ADMBA in situ hybridization (ISH) gene expression database onto age matched DevCCF templates to guide and validated the annotations (Fig. 3c; Extended Data Fig. 2). Moreover, we achieved segmentation and validation utilizing a multitude of additional evidence including contrast and autofluorescence features from co-registered DevCCF MRI and LSFM templates, and LSFM images with 3D histological labeling (e.g., fluorescence Nissl, neurofilament staining) (Fig. 3c; See methods for additional details).

**Figure 3.**
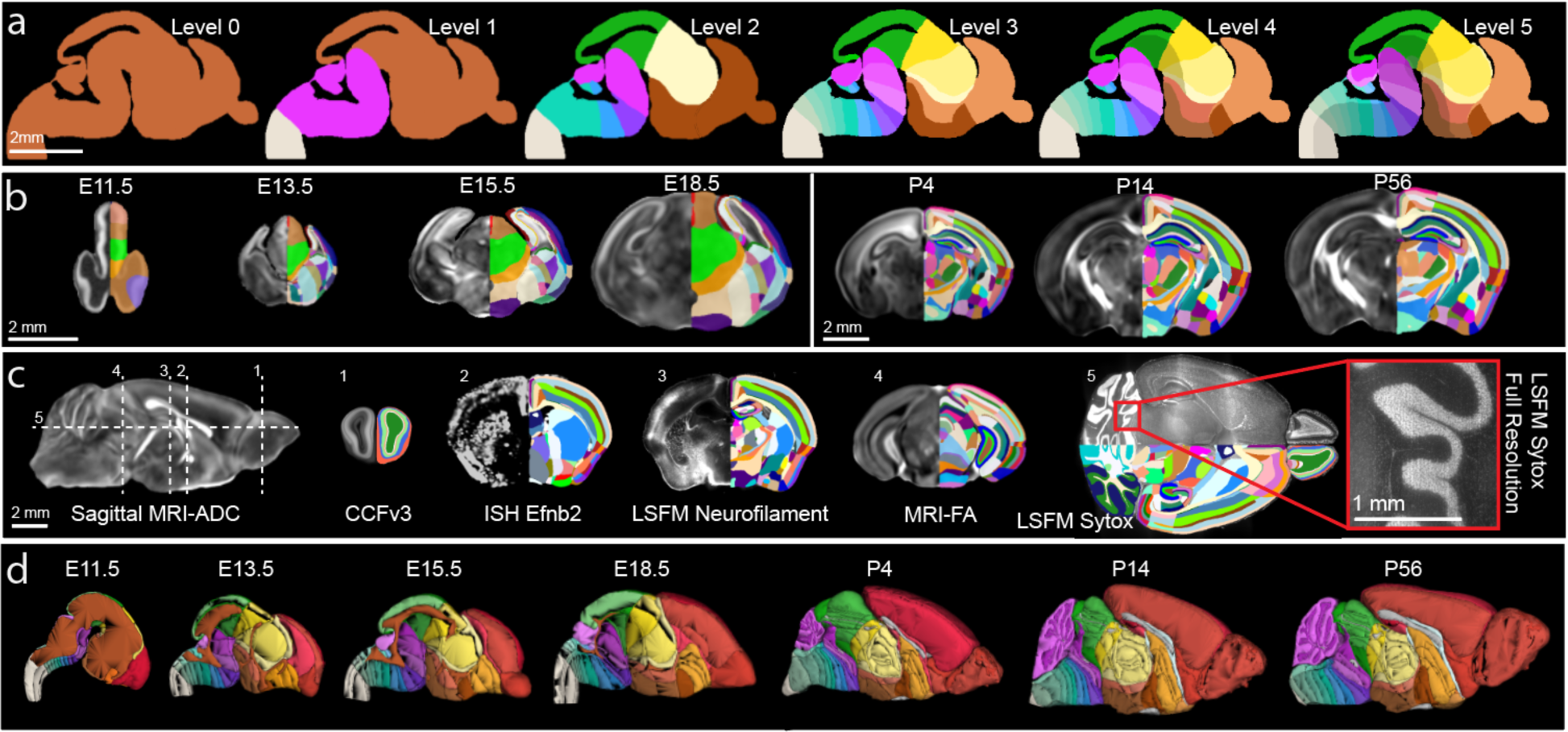
3D anatomical labels based on a developmental ontology. (a) DevCCF ontology levels 0 through 5 displayed over an E15.5 sagittal slice. (b) Coronal MRI FA slices through the subpallium with anatomical segmentation overlays on the right hemisphere. Embryonic and postnatal brains are depicted at uniform scales, respectively. (c) Gene expression, antibody and histological staining, different template contrasts registered to the DevCCF P56 to guide segmentations. On the left: midline sagittal MRI ADC slice indicating location of each of the following coronal (labels 1-4) and horizontal (label 5) slices. (1) CCFv3 template; (2) ISH Efnb2 gene expression; (3) LSFM neurofilament staining, (4) MRI FA; (5) LSFM SYTOX nucleus staining with densely packed cerebellar granule cells. (d) 3D renderings of DevCCF annotations, not to scale.

The DevCCF segmentations are simplest at E11.5 and grow in complexity through P56 (Fig. 3b,d). E11.5 annotations consist of ventricles, neuromeric boundaries in the rostral to caudal direction; floor, basal, alar, and roof boundaries in the ventral to dorsal direction; and pallium, subpallium divisions of the telencephalon (Fig. 3d). E13.5 annotations include additional subpallial segmentations as well as early stages of the concentric ring topology, defining early divisions of the neocortex, allocortex, and mesocortex (Fig. 3d). Cortical layers are segmented as early as P4, and cerebellar layers are segmented as early as P14 (Fig. 3b,d). Segmentations are mutually exclusive and comprehensively exhaustive, meaning all brain voxels are labeled uniquely with a single ontology ID.

### DevCCF allows examination of GABAergic neuron spatiotemporal development

The DevCCF offers new opportunities to map and quantify distinct cell types in developing brains using high resolution 3D microscopy, as done in adult mouse brains^7,39^. Here, we used Gad2-Cre;Ai14 mice that genetically label GABAergic neurons to examine their emergence in developing brains by applying tissue clearing methods and LSFM imaging at E11.5, E13.5, and E15.5 (Fig 4).

**Figure 4.**
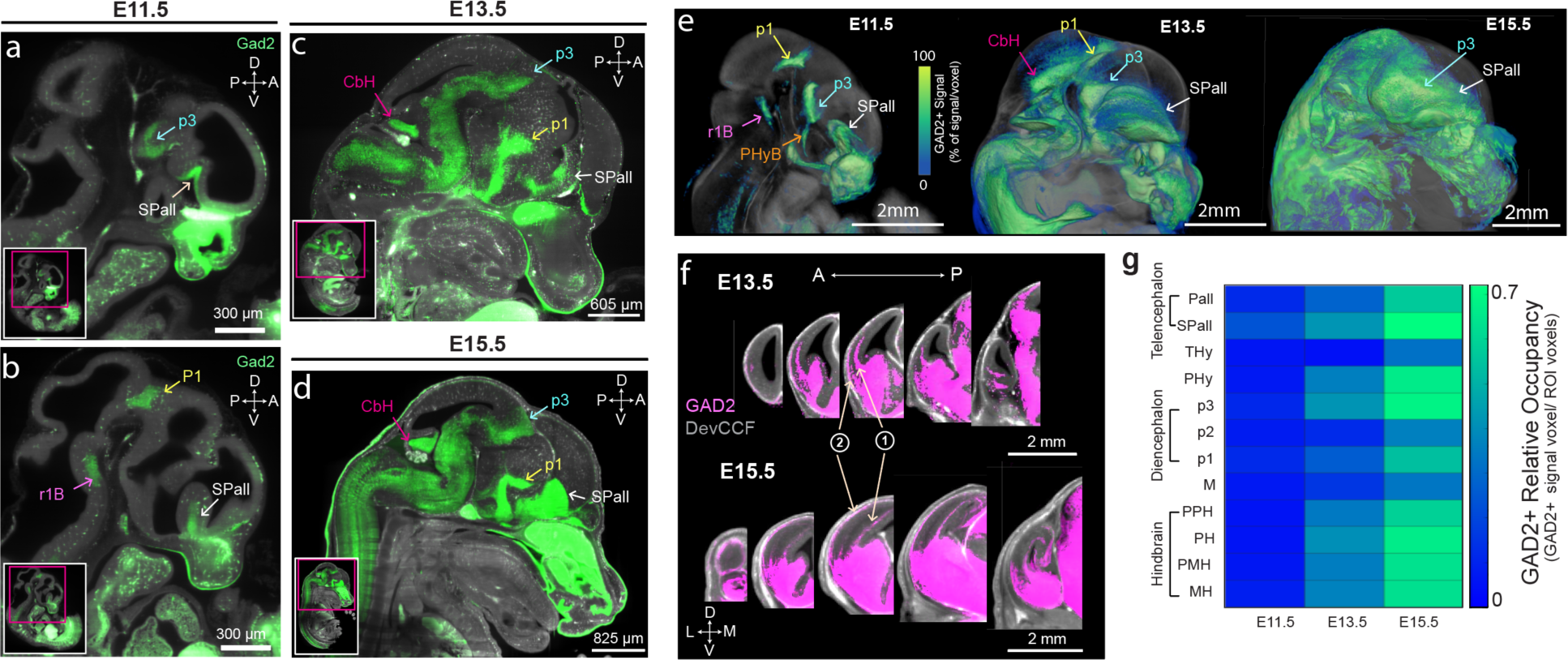
DevCCF to quantify early emergence of GABAergic neurons in embryonic brains. (a-b) Sagittal images of LSFM imaging from E11.5 Gad2-Cre;Ai14 mice show clusters of GAD2+ neurons in the subpallium (SPall), prosomere 1 and 3 (p1 and p3), rhombomere 1 basal plate (r1B). (c-d) The E13.5 (c) and E15.5 (d) brains show additional GAD2+ neurons in the cerebellar hemisphere (CbH). (e) Average GAD2+ signals at E11.5 (n=7), E13.5 (n=7), and E15.5 (n=5) with 3D rendering overlay on DevCCF templates. Note rapid expansion of GAD2+ neurons at E13.5 and E15.5 from initial clusters at E11.5. (f) In addition to local expansion, GAD2+ neurons migrate to deep (1) and superficial areas (2) of the pallium to establish cortical interneurons. (g) Quantification of GAD2+ signals in developmental neuromeres using DevCCF segmentations at E11.5, E13.5, and E15.5. Additional abbreviation: Pallium (Pall), terminal hypothalamus (THy), peduncular hypothalamus excluding telencephalon (PHy*), prosomere 2 (p2), midbrain (M), prepontine hindbrain (PPH), pontine hindbrain (PH), pontomedullary hindbrain (PMH), and medullary hindbrain (MH).

GABAergic neurons appear as focal clusters mostly in the subpallium (SPall), basal peduncular hypothalamus (PHyB), prosomere 1 (p1), prosomere 3 (p3), and rhombomere 1 basal plate (r1B) as a part of prepontine hindbrain (PPH) by E11.5 (Fig. 4a-e). Moreover, GABAergic neurons appear in the cerebellar hemisphere (CbH) by E13.5 with strong expression (Fig. 4c-e). In addition to local expansion of GABAergic neurons, this mapping clearly visualizes chains of migrating cortical interneurons from the SPall to the superficial and deep cortical areas (Fig. 4f). GAD2+ neurons continue to proliferate during embryonic development, occupying a greater relative volume of each brain region including their rapid expansion in the hindbrain (Fig. 4c-f). Using DevCCF segmentations, quantitative analysis revealed the spatiotemporal emergence of GABAergic neurons in developing brains (Fig. 4g).

### DevCCF enables CCFv3 aligned data analysis with a developmental framework

The P56 DevCCF provides a unique opportunity to compare developmental ontology-based labels with widely used CCFv3 annotations based on cytoarchitectures^15^. We mapped the CCFv3 labels to the stereotaxically aligned P56 DevCCF template (Fig. 5a)^14^. While the CCFv3 offers finer segmentation in the isocortex, DevCCF labels provide deeper segmentations in other areas such as the olfactory bulb, the hippocampus, and the cerebellum (Fig. 5a,b). Moreover, DevCCF isocortical regions have matching cortical layers with CCFv3, but major cortical areas are defined according to those of the ADMBA and the concentric ring topology in the rostrocaudal and mediolateral directions (Fig. 5a,b). We showed the spatial relationship between anatomical labels from each atlas and found several disagreements (Fig. 5b,c). For instance, prosomere 2 (p2) in the diencephalon of the DevCCF annotations overlaps with parts of the hypothalamus (HY), thalamus (TH), and midbrain (MB) in the CCFv3 annotations (Fig. 5b,c), highlighting differences in the segmentation criteria of the two labels. By integrating existing CCFv3 labels into DevCCF templates, users can easily decide their choice of anatomical labels to interpret signals of their interest in the same DevCCF template space. Further, the two annotations may be used in a common space to complement one another by functionally segmenting regions that are not annotated in either atlas individually, such as the layers of individual cerebellar lobules (Fig. 5b).

**Figure 5.**
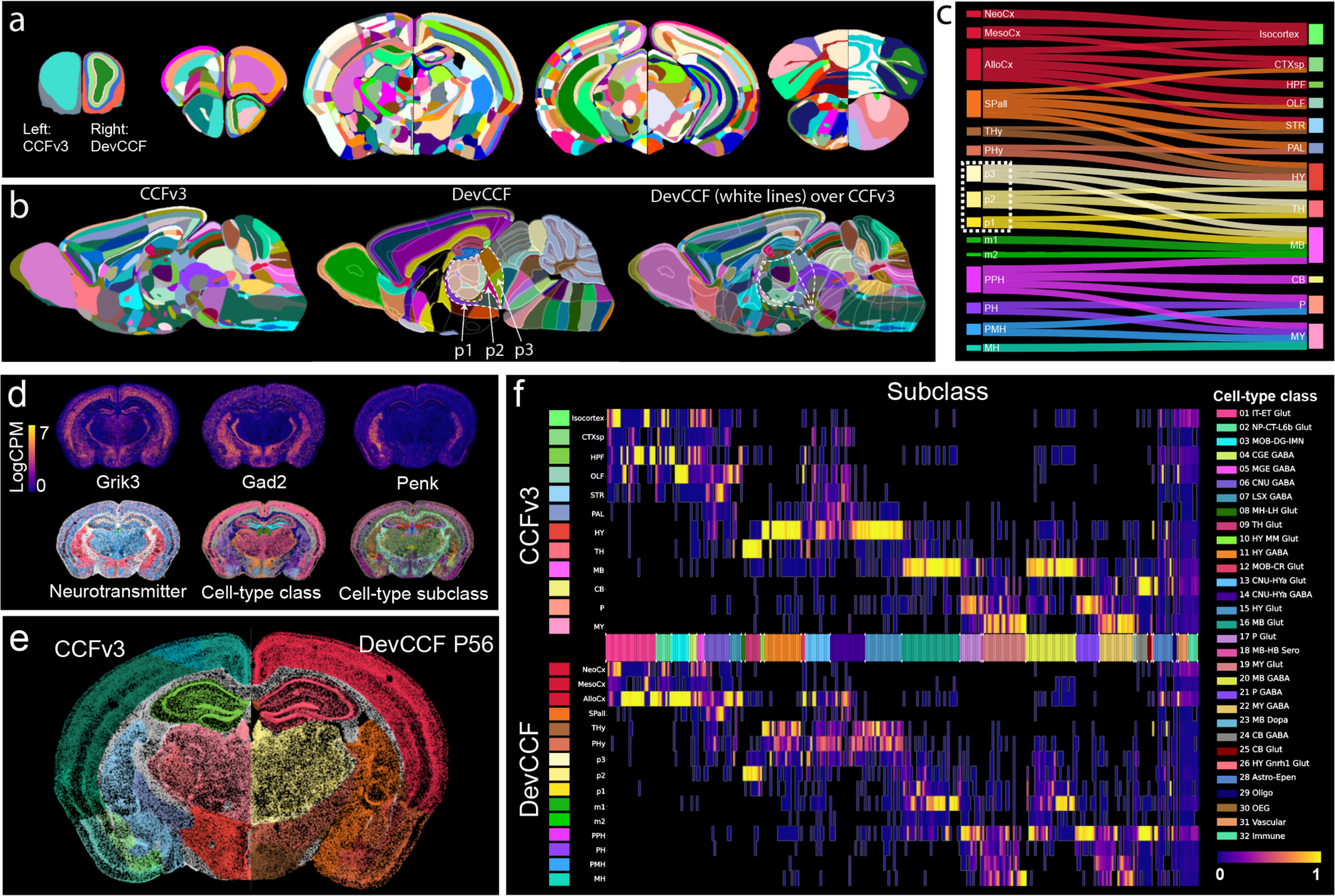
DevCCF enables CCFv3 aligned data analysis with developmental framework. (a) CCFv3 annotations (left hemisphere) and DevCCF P56 annotations (right hemisphere) overlayed on the DevCCF P56 template across five slices from anterior (left) to posterior (right). (b) Sagittal slices with CCFv3 annotations (left), DevCCF (middle), and DevCCF label boundaries overlaid on CCFv3 annotations (right). Neuromeric boundaries of the prosomere (p)1, p2, p3 highlighted with white dashed lines over colored CCFv3 annotations demonstrate shared and non-overlapping boundaries of CCFv3 and DevCCF annotations. (c) Sankey diagram illustrates matched structural relationship between DevCCF and CCFv3 annotations. Size of individual areas represents logarithmic scale of regional volume. (d) Single section of MERFISH spatial transcriptome data with three representative genes (top) and cell type classifications (bottom). (e) Registered spatial transcriptome with CCFv3 segmentation (left) and DevCCF segmentation (right). Colors are the same as in (c). (f) Heatmap of cell-type distribution in CCFv3 (top) and DevCCF segmentations (bottom). Heatmap values show proportion of cells in a region relative to the total number of cells in the subclass. Cell-types are ordered by their parent class, denoted by color bars on the x-axis.

Leveraging co-registered DevCCF and Allen CCFv3 labels, one can easily assess data previously mapped to the CCFv3 in a developmental context (Fig. 5d-f). For instance, a study using a spatial transcriptome approach based on Multiplexed Error-Robust Fluorescence in situ Hybridization (MERFISH) identified 32 classes and 306 subclasses of cell types across the whole mouse brain based on CCFv3 ontology (Fig. 5d)^40^. We re-mapped the cell type classification data in the DevCCF and compared the distribution of cell types in the two atlas labels. Mirroring segmentation differences (Fig. 5b,c), cell-type class and subclass distribute differently based on DevCCF labels with developmental ontology. For example, cell types enriched in the CCFv3 hypothalamus (HY) segregated mostly into the DevCCF terminal hypothalamus (THy), peduncular hypothalamus (PHy), and prosomere 3 (p3), highlighting the potential contribution of different cell types in areas with distinct developmental ontology (Fig. 5f). Together, seamless re-interpretation of CCFv3 mapped cell type data using DevCCF can facilitate the interoperability and data integration across different studies.

### Web visualization for DevCCF

Neuroglancer based web visualizations (https://kimlab.io/brain-map/DevCCF/) allow users to interactively explore the DevCCF in 3D. The web visualizations feature all co-registered MRI and LSFM templates as well as DevCCF annotations at the seven developmental ages. The P56 DevCCF also includes co-registered CCFv3 annotations to compare with DevCCF annotations. Neuroglancer webpages include 2D slice viewers in 3 orthogonal planes that may be resliced to view in any orientation. Each template and annotation dataset may be selected as an independent layer and viewed and edited with user defined color, contrast, and opacity (Fig. 6a,b). Hovering over a region reveals its name, abbreviation, and assigned numerical identifier. Region selection reveals additional information, such as region abbreviation, subregions, and parent regions in the ontology tool (Fig 6c). Users can apply these interactive tools to create layer structures that highlight their desired anatomical features in a single template (Fig. 6d-f) or explore features of multiple templates at once to gain additional anatomical context in a single viewer (Fig. 6g-i). All DevCCF templates and annotations are freely available for download (see data availability section).

**Figure 6.**
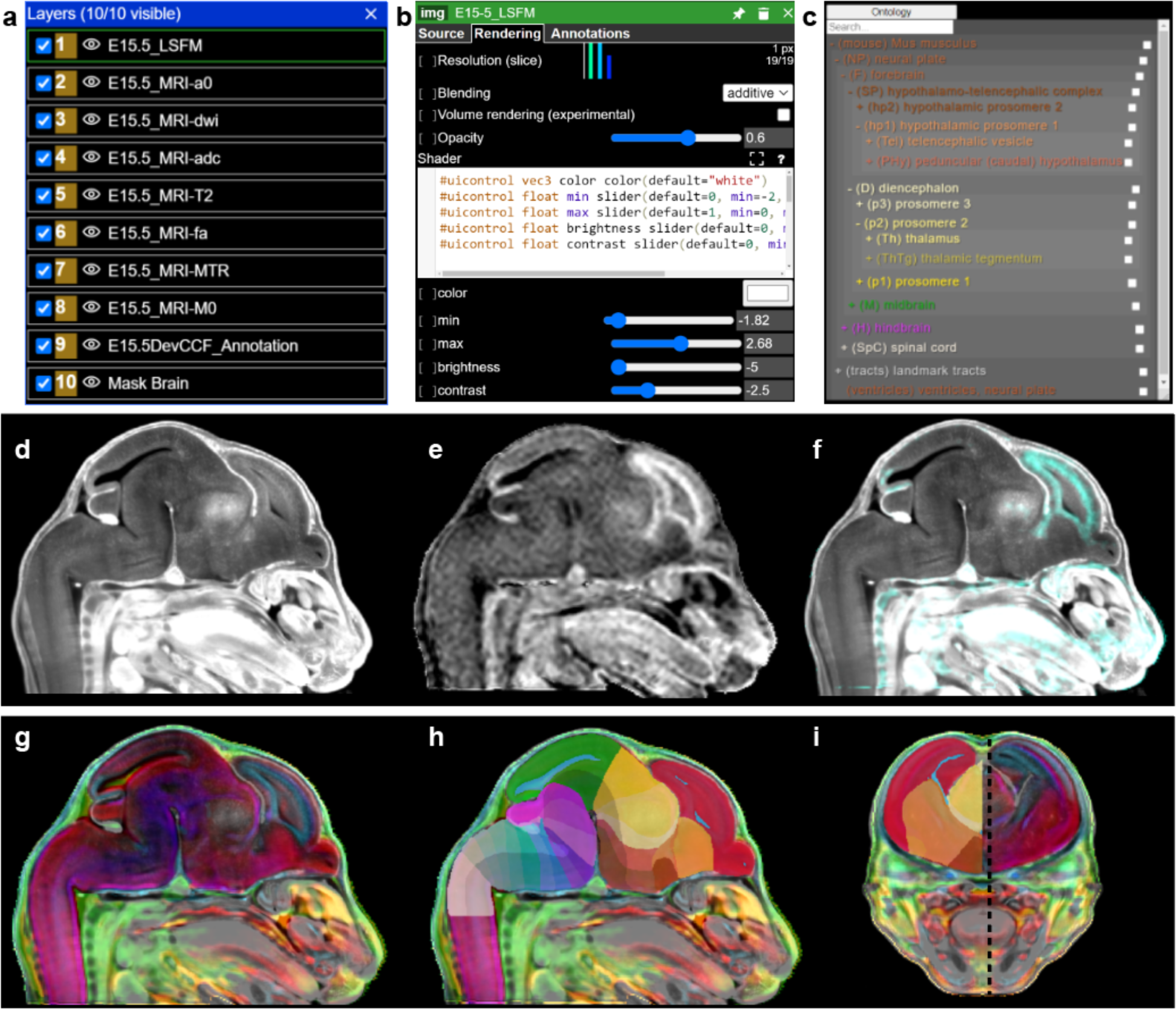
Web Visualization for DevCCF. (a) Layer panel allows layer selection with a right click and hiding by selecting the eye icon to the left of the layer. (b) Once selected, the layer edit tool enables user modification of layer color, contrast, and brightness. (c) The ontology viewer allows search and selection of individual segmentations and parent regions. When a region is selected in the viewer or ontology tool, the region’s metadata is displayed including region name, abbreviation, and ID. The ontology tool may be dragged to any location in the viewer. (d-f) Neuroglancer allows users to visualize either individual E15.5 LSFM autofluorescence in (d), MRI MTR template in (e), or their overlay (f). (g-i) E15.5 DevCCF templates overlaid (g) and segmentation (h, i). Black dashed line in (i) denotes sagittal slice location of (g,h).

## Discussion

Here, we present the DevCCF as a set of 3D atlases for the developing mouse brain including MRI and LSFM templates and corresponding 3D annotations based on a developmental ontology. The DevCCF can facilitate interpretation and integration of diverse data across modalities and scales including advanced high resolution 3D imaging and spatial genome data^41^.

The DevCCF contains undistorted morphology and intensity averaged symmetric templates at seven key developmental ages: E11.5, E13.5, E15.5, E18.5, P4, P14, and P56. Template ages correspond with the ADMBA and cover critical stages of regional differentiation^24^. While individual brains have unique differences, morphologically averaged templates from male and female samples provide a common reference morphology at each age. Furthermore, the stereotaxically aligned postnatal templates provide spatial coordinates to guide surgical procedures (e.g., stereotaxic brain injection). At each age, the DevCCF contains at least four MRI contrast templates along with an aligned age-matched LSFM autofluorescence template, which helps to register data from different modalities^20,42,43^. For instance, we mapped whole brain spatiotemporal trajectories of GABAergic neurons from LSFM imaging, spatial transcriptome, and ADMBA ISH data to various DevCCF ages, demonstrating how researchers can easily align data from different imaging methods to the DevCCF. Hence, DevCCF templates enable precise spatial localization of data and promote comparative analyses across diverse datasets through development.

The DevCCF 3D annotations are adapted from 2D ADMBA labels based on the prosomeric model of mammalian brain development^24,28^. These annotations fit the current understanding of developmental anatomy with the notable re-organization of the pallium into a concentric ring topology^35,38^. The expert guided 3D digital annotations allow users to examine labels with any desired angle and serve as a neuroinformatic tool to automatically quantify signals across different brain areas in the developing mouse brain^15,44–46^.

Inconsistencies of anatomical segmentation between different atlases have caused confusion within the community^14^. We also observed partially overlapping, yet distinct borders with varying depth of segmentations between community standard CCFv3 and DevCCF based on two different delineation criteria. Therefore, we aligned the CCFv3 template and annotations to DevCCF postnatal templates to place the two atlas annotations in the same spatial template. This allows datasets previously registered to the CCFv3 to seamlessly integrate with the DevCCF with undistorted stereotaxic spatial coordinates and developmental annotations. This atlas integration will promote large scale data analysis to accommodate rapidly growing cell census data^47^.

We envision that the DevCCF will be widely used to support various mouse brain mapping experiments. Advances in tissue clearing and high resolution microscopy have been critical to increase our ability to examine individual cell types in the whole mouse brain at single cell resolution^18,46,48,49^. MRI has been used to examine macroscopic changes in brain morphology, connectivity, and function^25,50–55^. Furthermore, recently developed spatial transcriptomic approaches have begun to unravel the gene expression landscape in the adult and developing mouse brain^8,10,11,40,56–59^. Much of this new data can be mapped onto the DevCCF at matched ages for analysis to enhance interoperability of different studies. However, the DevCCF is not without limitations. Adding more samples at each age can help improve template quality^15^.

Additionally, the inclusion of other 3D imaging modalities such as micro-computed tomography could expand the DevCCF’s utility. As advancements in multimodal registration techniques progress, alignment between different modalities may benefit from enhanced registration methods^42,43,60,61^. Moreover, the selected seven developmental ages, though comprehensive, may be too sparse to fully capture the rapid cellular changes occurring in early brain development.

Cross-age registration and interpolation can generate mappings between template ages that characterize morphological changes and examine aligned datasets over the temporal axis^25^. We also expect that ongoing discoveries will necessitate further segmentation and updates to DevCCF annotations. Emerging artificial intelligent algorithms can drive automated segmentation based on diverse mapped data^62,63^. Importantly, grounded in the prosomeric model of vertebrate development, the DevCCF can serve as the foundation for constructing developmental atlases for other mammalian species (e.g., macaque, human)^64–66^. Therefore, future efforts should focus on continued refinement of the DevCCF and analogous atlases in diverse species.

In conclusion, the DevCCF establishes a standardized anatomical framework for investigating developing mouse brains and facilitates collaborative and reproducible advancements in neuroscience research.

## Methods

### Animals

All experiments and techniques involving live animals have been approved and conform to the regulatory standards set by the Institutional Animal Care and Use Committee (IACUC) at the Pennsylvania State University College of Medicine. We used in-house bred C57bl/6J mice (originally purchased from the Jackson Laboratory, Strain #:000664) or transgenic animals with C57bl/6J background to create MRI and LSFM templates. For brain-wide labeling of pan-GABAergic cell types during embryonic and early postnatal development, we used Gad2-IRES-Cre mice (The Jackson Laboratory, stock 028867) crossed with Ai14 mice which express a Cre-dependent tdTomato fluorescent reporter (The Jackson Laboratory, stock 007908). We used tail samples with PCR for genotyping including Rbm31-based sex genotyping for mice younger than P6. All mice were maintained under a 12-hour light/12-hour dark cycle at 22–25 °C with access to food and water ad libitum.

### Timed pregnancies and brain sample collection

Timed pregnancies followed recommendations from The Jackson Laboratory (https://www.jax.org/news-and-insights/jax-blog/2014/september/six-steps-for-setting-up-timed-pregnant-mice). Adult breeder males were singly housed for 1 week prior to pairing. We paired a male and a female breeder in the evening, removed the male breeder the following morning, and checked for the presence of a vaginal plug in the females. After separation, we measured the baseline body weight of all females on what we considered E0.5 and added extra enrichment to the cages for female breeders. Subsequently, we weighed the females again on E7.5 and E14.5 to assess weight gain, expecting a 1 g or 2-8 g gain at these respective timepoints. Pregnancy was confirmed either by the presence of fetuses during euthanasia of the dams and fetal tissue collection (on E11.5, E13.5, E15.5, and E18.5) or by the birth of a litter.

On target collection dates for embryonic samples (E11.5, E13.5, E15.5, and E18.5), pregnant dams were placed in an isoflurane chamber until deeply anesthetized at which point a mixture of ketamine and xylazine was administered via intraperitoneal injection. The dams were subjected to cervical decapitation once fully anesthetized. Then, the uterine horns were immediately removed and placed in ice-cold petri dishes filled with 0.05 M PBS for careful removal of embryos from the uterine casing. E18.5 brains were dissected out at this point while E11.5, E13.5, E15.5 were processed as whole embryos. Embryonic samples were incubated in a 4% paraformaldehyde (PFA) in 0.05 M PBS solution for two days at 4°C before storing in 0.05 M PBS at 4°C until use. All embryos were characterized according to the Theiler Staging (TS) Criteria for Mouse Embryo Development^67^. We used E11.5 for TS19, E13.5 for TS21, E15.5 for TS24, and E18.5 for TS26. After PFA fixation was complete, dissection of the embryos from the yolk sac (E11.5) and eye/ eye pigment removal (E11.5, E13.5, E15.5) were performed under a dissection microscope (M165 FC Stereomicroscope, Leica), followed by storage in 0.05 M PBS at 4°C until use. Whole embryos were used for E11.5, E13.5, and E15.5, and dissected brains were used for E18.5 for tissue clearing and LSFM imaging. All embryonic samples were tailed for sex genotyping.

For postnatal brains (P4, P14, and P56), we defined pups at birth as P0. For collection, mice were anesthetized by a mixture of ketamine and xylazine via intraperitoneal injection. Anesthetized animals were subsequently perfused by saline (0.9% NaCl) and freshly made 4% PFA. Decapitated heads were fixed in 4% PFA at 4°C overnight, followed by brain dissection and storage in 0.05 M PBS at 4°C until use.

The Kim Lab at Penn State University prepared all animal samples and performed tissue clearing with LSFM imaging. Whole embryos (E11.5, E13.5, E15.5) or decapitated samples (E18.5, P4, P14, and P56) were sent to Dr. Jiangyang Zhang’s lab at NYU for high resolution MRI.

### Magnetic Resonance Imaging

All imaging was performed on a horizontal 7 Tesla MRI system (Bruker Biospin, Billerica, MA, USA) equipped with a high-performance gradient system (maximum gradient strength of 670 mT/m). We used a transmit volume coil (72 mm inner diameter) together with a 4-channel receive-only phased array cryoprobe with high sensitivity. Hair and scalp were removed and heads with intact skulls were imaged to prevent deformations. As the embryonic mouse has immature soft skulls, embryonic mouse heads were embedded in 5% agarose gel (Sigma Aldrich, St Louis, MO, USA) for additional support. Specimens were placed in 5 mL syringes filled with Fomblin (Solvay Solexis, Thorofare, NJ, USA) to prevent dehydration and susceptibility to artifacts.

High-resolution diffusion MRI was acquired using an in-house 3D diffusion-weighted gradient and spin-echo (DW-GRASE) sequence^68^ with the following parameters: echo time (TE)/repetition time (TR) = 30/400 ms, two signal averages, diffusion gradient duration/separation = 4/12 ms, 60 diffusion directions with a b-value of 1.0 ms/µm^2^ for E11.5, 2.0 ms/µm^2^ for E13.5-E17.5, and 5.0 ms/µm^2^ for P4, P14, and P56 brains. The increase in b-values with age was necessary as the diffusivity of brain tissues decreases with development^69^. Co-registered T2-weighted data were acquired using the same sequence but with TE/TR = 50/1000 ms. Co-registered magnetization transfer-weighted (MT) and reference data (M0) were acquired using the same sequence with TE/TR = 8/700 ms, and the following MT parameters: offset frequency= 5kHz, 100 Gaussian pulses with a duration of 3 ms and an amplitude of 8 µT. The native and interpolated spatial resolutions of the MRI data were 0.063/0.0315 mm for E11.5, 0.068/0.034 mm for E13.5, 0.075/0.037.5 mm for E15.5, 0.08/0.04 mm for E18.5, and 0.1/0.05 mm isotropic for P4-P56.

The 3D MRI data were reconstructed from k-space to images and zero-padded to twice the raw image resolution in each dimension in MATLAB (Mathworks, Natick, MA, USA). Magnetization transfer ratio map was computed using MTR = 1-MT/M0. Diffusion tensor images^70^ were constructed using the log-linear fitting method in DTI Studio (http://www.mristudio.org), and the tensor-based scalar metrics were generated, including the mean diffusivity (MD) and fractional anisotropy (FA).

### Tissue Clearing

We mainly used SHIELD (Stabilization under Harsh conditions via Intramolecular Epoxide Linkages to prevent Degradation) tissue clearing to ensure minimal tissue volume changes while preserving endogenous fluorescence signals when available^71^. Commercially available SHIELD preservation, passive clearing reagents, and detailed protocols were obtained from LifeCanvas Technologies (https://lifecanvastech.com/). For P56 brains, PFA-fixed samples were incubated in SHIELD OFF solution for 4 days at 4°C on an orbital shaker. Subsequently, the SHIELD OFF solution was replaced with SHIELD ON buffer and incubated for 24 hours at 37°C with shaking. Tissues were incubated in 20 mL of delipidation buffer at 37°C for 4-6 days followed by an overnight wash in 1x PBS at 37°C with gentle shaking. If sample imaging did not occur within a week after the delipidation step, tissue samples were stored in 1x PBS containing 0.02% sodium azide at 4°C before continuing to the next step. To match the refractive index (RI) of the delipidated tissues (RI= 1.52) and obtain optical clearing, samples were incubated in 20 mL of 50% EasyIndex + 50% distilled water for 1 day, then switched to 100% EasyIndex solution for another day at 37°C with gentle shaking. For samples in earlier time points, we used whole embryos (E11.5, E13.5, and E15.5) or dissected brains (E18.5, P4 and P14) with the same protocol but with smaller reagent quantities and shorter incubation times. Once cleared, embryonic samples were stored in tightly sealed containers with 100% EasyIndex at room temperature (20-22°C). For 3D immunolabeling and histological staining, we used electrophoresis based active clearing using the SmartClear II Pro (LifeCanvas) and active labeling using SmartLabel (LifeCanvas) based on protocols provided by LifeCanvas. Briefly, after the SHIELD OFF step described above, samples were incubated in 100 mL of delipidation buffer overnight at room temperature (RT). Samples were inserted in a mesh bag, placed in the SmartClear II Pro chamber, and delipidated overnight. Samples were transferred to 20 mL of primary sample buffer and incubated overnight at RT. Samples were placed in a sample cup in the SmartLabel device with an antibody cocktail to perform active labeling^72^. We used neurofilament H (Encor, cat. no. MCA-9B12, 10 µl per hemisphere) with Alexa Fluor 647 anti-mouse IgG secondary antibody (Jackson Immuno Research, cat. no.: 715-607-003, RRID: AB_2340867, 2.7 µl per hemisphere), and propidium iodide (Thermo Fisher, cat. no.: P1304MP, 12 µl /hemisphere) for pan-cellular labeling. For the limited dataset at P56, we used a modified iDISCO-based tissue clearing method as previously described^73^.

### Light Sheet Fluorescence Microscopy Imaging and 3D reconstruction

For LSFM imaging, all samples were embedded in an agarose solution containing 2% low-melting agarose (Millipore Sigma, cat. no.: A6013, CAS Number: 9012-36-6) in EasyIndex using a custom sample holder. Embedded samples were then incubated in EasyIndex at room temperature (20-22°C) for at least 12 hours before imaging using the SmartSPIM light sheet fluorescence microscope (LifeCanvas). During the imaging process, the sample holder arm securing the embedded sample was immersed in 100% EasyIndex. Our imaging setup consisted of a 3.6X objective lens (LifeCanvas, 0.2 NA, 12 mm working distance, 1.8 μm lateral resolution), three lasers with wavelengths of 488 nm, 560 nm, and 642 nm, and a 5 μm z step size. After imaging, all samples were stored in 100% EasyIndex at room temperature (20-22°C). For our iDISCO cleared P56 brain samples, we did not use agarose embedding and directly mounted samples in a custom-built holder. LSFM imaging was performed in ethyl cinnamate for index matching (Millipore Sigma, cat.no.: 112372, CAS number: 103-36-6) using the same imaging parameters.

For 3D reconstruction, we developed a parallelized stitching algorithm optimized for conserving hard drive space and memory consumption initially based on Wobbly Stitcher^17^. The algorithm started by collecting 10% outer edges of each image tile and making a maximum intensity projection (MIP) of outer edges in the axial (z) direction for every set of 32 slices of the entire stack. The algorithm then aligned z coordinates of MIP images across image columns, followed by the x and y coordinate alignment. Finally, 32 slices within each MIP were adjusted based on curve fitting to reach final coordinates of each tile. This algorithm only reads the raw images two times (at the beginning and the final writing), which significantly reduced the bottleneck of reading large files in a storage drive.

### Symmetric Template Construction

Each symmetric template is an intensity and morphological average of multiple male and female samples with a sample size ranging from 6 to 14 (Extended Data Table 1). After stitching, images were preprocessed for template construction. MRI data preprocessing involved (1) digital postnatal brain extraction and (2) sample orientation correction. LSFM data preprocessing involved (1) image resampling to 3 sizes: 50 µm, 20 µm, and 10 µm isotropic voxel resolution, and (2) sample orientation correction to ensure all images were facing the same direction. To ensure template symmetry, each preprocessed image was duplicated and reflected across the sagittal midline, doubling the number of input datasets used in the template construction pipeline. Template construction, using functionality contained in ANTs^34,74^, was employed on Penn State’s High-Performance Computing system (HPC). Briefly, starting from an initial template estimate derived as the average image of the input cohort, this function iteratively performed three steps: (1) non-linearly registered each input image to the current estimate of the template, (2) voxel-wise averaged the warped images, and (3) applied the average transform to the resulting image from step 2 to update the morphology of the current estimate of the template. Iterations continued until the template shape and intensity values stabilized. MRI templates were constructed at their imaged resolution using ADC MRI contrasts for initial postnatal templates and diffusion weighted imaging (DWI) contrasts for embryonic templates. Once the initial MRI template was constructed, the sample to template warp fields were applied to all MRI contrasts for each sample. Warped samples were averaged to construct templates for each contrast. LSFM templates were constructed from autofluorescence data collected from C57bl/6J mice and transgenic mice with a C57bl/6J background. To save memory and improve speed, LSFM templates were initially constructed at 50 µm isotropic resolution. This template was resampled for template construction initialization at 20 µm isotropic resolution, a process repeated to construct the final LSFM template with 10 µm isotropic resolution input images.

### Multimodal registration of 3D imaging to the DevCCF

We aligned the CCFv3 to the P56 DevCCF and each LSFM template to the corresponding age matched DevCCF MRI template to enable data integration across both modalities with undistorted morphology. Our protocol aims to address multimodal registration challenges due to differences in brain and ventricle volume that often result in internal structure misalignment.^75^ We performed initial non-linear registration of the 3D datasets (CCFv3 and LSFM templates) to the age-matched DevCCF MRI template using ANTs with the mutual information similarity metric^34^. We then visually compared the warped 3D dataset with the DevCCF template in ITK-SNAP^76^ to identify landmark brain regions that remained misaligned after the initial registration. Whole brain masks and misaligned regions were segmented for the 3D dataset and DevCCF template in 3D using Avizo (Thermo Fisher Scientific). The segmented regions were subtracted from the brain masks, creating modified brain masks with identifiable boundaries around misaligned brain regions. This provided a map of regions that needed correction. Next, linear registration was performed of the modified brain masks, followed by equally weighted non-linear alignment of both the 3D data images and their modified brain masks. Landmark-assisted multimodal registration warp fields were resampled and applied to transform CCFv3 and LSFM templates to MRI template morphology at 20 μm isotropic voxel resolution. CCFv3 annotations were also transformed to MRI template morphology at 20 μm isotropic voxel resolution for comparison with the DevCCF. We also registered LSFM whole brain data featuring immunostaining or Cre-dependent fluorescence to the DevCCF to assist with annotation, segmentation, and validation. Each LSFM dataset collected up to three channels simultaneously, always including one autofluorescence channel. We used non-linear registration to align LSFM autofluorescence channel data to the DevCCF LSFM template. The LSFM template was chosen to achieve optimal registration quality due to matching the autofluorescence contrasts. Forward transforms were applied to LSFM immunostaining and Cre-dependent fluorescence channel data to align them to the DevCCF template. To re-interpret Allen Institute MERFISH data previously aligned to the CCFv3 with the DevCCF label, we (1) used the inverse transforms to warp the DevCCF annotations to the CCFv3 template, then (2) applied the DevCCF annotations to the MERFISH dataset.

### 2D in situ hybridization gene expression mapping onto the DevCCF

We downloaded in situ hybridization (ISH) data consisting of 2D images of coronal and sagittal sections stained for gene expression from the ADMBA at each age^24^. We warped the ISH data to the respective MRI templates using ANTs^34^ with the following steps: (1) Sample level data was compiled across ISH experiments, intensity inverted, and resampled to 512×512 pixels for computational tractability. (2) Sample slices were reconstructed to a 3D volume and empty slices were generated using a B-spline scattered data approximation technique^77^. (3) The MRI template was aligned to the sample reconstruction using linear registration. (4) Slice-wise non-linear registration was performed on each ISH slice to the corresponding MRI slice using a mutual information metric to improve within subject consistency. (5) The MRI template was aligned to the slice-wise corrected sample reconstruction using nonlinear registration. (6) The inverse transform from step 5 was used to transform the ISH sample to the DevCCF template. (7) ISH slice-wise corrected sample reconstruction was divided slice by slice to individual gene expression patterns, empty slices were generated with B-spline scattered data approximation, and individual gene reconstructions were transformed to the DevCCF morphology.

### Anatomical segmentations for the DevCCF

We performed theory and data-driven anatomical segmentation of the multimodal templates at each age using Avizo (Thermo Fisher Scientific). We manually drew contours on coronal, horizontal, and sagittal slices of templates to generate 3D segmentations. To assist in the process, we used various interpolation, thresholding, and smoothing tools. We assigned unique labels and colors to each region and further developed the hierarchical nomenclatures following standards in the ADMBA.

We followed the principles of the prosomeric model to define brain regions based on morphological features, such as sulci, fissures, ventricles, commissures, tracts, cytoarchitecture, and gene expression^28^. Segmentations started with large regions defined early in development, such as the neural plate, and were progressively subdivided into smaller regions as defined by the prosomeric model, such as the forebrain, hindbrain, and spinal cord until reaching a level of detail comparable to at least level 5 (fundamental caudo-rostral and dorso-ventral partitions) in all brain regions. Additionally, we updated the telencephalon to reflect the cortical concentric ring topology^35^. Anatomical divisions and subdivisions were drawn across the whole brain in multiple segments and combined. For example, floor, basal, alar, and roof plate segmentations were drawn separately from neuromere segmentations and later overlayed with one another. Similarly, cortical region segmentations (e.g., hippocampal cortex and olfactory cortex) were drawn separately from cortical layer segmentations (e.g., ventricular zone, mantle zone), and later overlayed to combine. This technique allows efficient segmentation correction upon validation. Additional brain regions were modified from the ontology described in the ADMBA to meet the current anatomical understanding^35,38,78^.

The morphological foundation of the DevCCF was primarily guided by MRI templates, additional details from LSFM templates and registered LSFM data. Here we provide a few examples of each data type. The MRI T2-weighted templates, in conjunction with LSFM templates, facilitated delineation of brain tissue from non-brain structures, as well as ventricle identification in embryonic DevCCF templates. MRI FA templates highlight white matter tract landmarks, which serve as markers for many boundaries (e.g., the boundary of p1 and p2 is immediately caudal to the retroflex tract). LSFM templates were instrumental in segmenting features such as the choroid plexus in the developmental roof plate. We imaged and aligned additional 3D fluorescent cell-type specific datasets to provide reference data for DevCCF segmentations. For example, we used LSFM images of embryonic GAD2-Cre;Ai14 samples to delineate the subpallium (SPall), prosomere 1 (p1), prosomere 3 (p3), and the cerebellum. Additionally, LSFM imaging of SYTOX stained mouse brains samples provided Nissl-like 3D histology, allowing us to use classical neuron cell density patterns during segmentation.

The ADMBA and associated ISH data were key resources to segment large rostro-caudal, and dorso-ventral boundaries. The atlas and gene expression data provided extensive supportive evidence for the prosomeric model of vertebrate brain development, aiding in validating relative relationships among neuromeres, dorso-ventral plates, nuclei, white matter tracts, and cranial nerve roots. Moreover, we utilized registered ISH data associated with the ADMBA to validate DevCCF segmentations.

The CCFv3^15^ provided delineations of the cortical layers and many nuclei in the P56 mouse brain. The CCFv3 template and annotations were aligned to the P56 MRI template using our landmark-assisted multimodal registration methods. Previously validated CCFv3 cortex layers (1, 2/3, 4, 5, 6a, and 6b) were imported to the P56 DevCCF. P56 DevCCF cortical regions (e.g., insular cortex, entorhinal cortex) were segmented manually and combined with imported CCFv3 layers to finalize cortical segmentations. Additional CCFv3 segmentations with corresponding prosomeric model regions (e.g., subdivisions of prosomere 2 and mesomere 1) were aligned to the DevCCF and used as primers to segment the DevCCF. CCFv3 cortical layers were also used for P14 and P56. CCFv3 cortical layers were similarly aligned from the P56 DevCCF to the P14 and P4 DevCCF templates via landmark-assisted multimodal registration methods and used to prime segmentation.

Anatomical segmentations were paired with numerical identifiers (IDs). IDs below 18000 represent previously existing ADMBA labels with potential minor name, abbreviation, and ontology level updates. Newly added structures to the DevCCF ontology have IDs in the range 18000-19999. To ensure compatibility with viewing and analysis tools requiring 16-bit depth labels, existing 32-bit ADMBA IDs that fell outside the 16-bit depth range (0-65535) were fit into the 20000 to 29999 range. This was achieved through a formulaic approach where the last four digits of the original 32-bit IDs were extracted and given a prefix of 2. New 16-bit labels ensure the modified IDs retain their uniqueness and do not overlap with either original or newly generated labels. Annotation names, abbreviations, parent structures, associated colors, previous 32-bit IDs, and current 16-bit IDs are organized in Supplementary Table 1.

### GABAergic neuron Quantification

We performed LSFM to image Gad2-Cre;Ai14 mouse embryos at E11.5, E13.5, and E15.5. We used ilastik^79^ to train a pixel classification-based machine learning model for each age to identify GAD2 positive (GAD2+) voxels in the whole embryo at 1.8 × 1.8 × 5 µm^3^ voxel resolution. We resampled images to 20 µm isotropic resolution for image registration to the DevCCF, where each voxel value represented the sum of GAD2+ voxels in the respective full resolution image. The GAD2+ voxel count was registered to age matched DevCCF templates as described above. GAD2+ voxel count relative occupancy per anatomical region was calculated as a ratio of the sum of positive voxels to the number of voxels per region.

### Key Resource List: See Extended Data Table 2

### Resource Availability

### Lead Contact

Additional information is available by contacting the corresponding author, Yongsoo Kim.

### Data & Code Availability

The Developmental Common Coordinate Framework (DevCCF) is an openly accessible resource. The DevCCF templates, labels, and additional documentation can be viewed and downloaded via the DevCCF hub at https://kimlab.io/brain-map/DevCCF/ or direct download link at DevCCF_MRI_sharable_v3.6. The DevCCF data viewer was built using Neuroglancer (https://github.com/google/neuroglancer). Additional data will be made available via https://kimlab.io as they are developed.

All available data will be deposited in public data repository (e.g., Mendeley data, GitHub) upon publication.

## Acknowledgement

We express gratitude to Yongsoo Kim Lab members and other DevCCF team members for their commitment, expertise, and motivation. We acknowledge the invaluable support of The Penn State College of Medicine High Performance Computing cluster. Our thanks to BioRender.com for their illustration generation platform. We are grateful to the members of the BRAIN Initiative Cell Census Network for their insights.

This work was supported by National Institutes of Health grants RF1MH12460501, R01NS108407, R01MH116176 (to Y.K.), and R01EB031722 (to J.G.).

The contents are solely the responsibility of the authors and do not necessarily represent the views of the funding agency.

## Author contribution

Y.K. and L.N. conceptualized the project; R.B., J.K.L., J.A.M., D.S., S.B.M. acquired LSFM data; J.Z. and C.H.L. acquired MRI data; Y.T.W. developed stitching pipeline; F.A.K., N.T., and J.G. performed template construction and image registration; F.A.K., R.P., and L.P. delineated anatomical segmentations; J.K.L. trained GABAergic machine learning model; A.B., and L.N. performed MERFISH Integration; J.T.D. and L.N. imported Allen developmental gene expression database; D.J.V. designed web visualization; Y.K., L.P., J.G., J.Z., and L.N. provided supervision; F.A.K. and Y.K. prepared the manuscript; All authors discussed and commented on the manuscript.

---

**Extended Data Figure 1.**
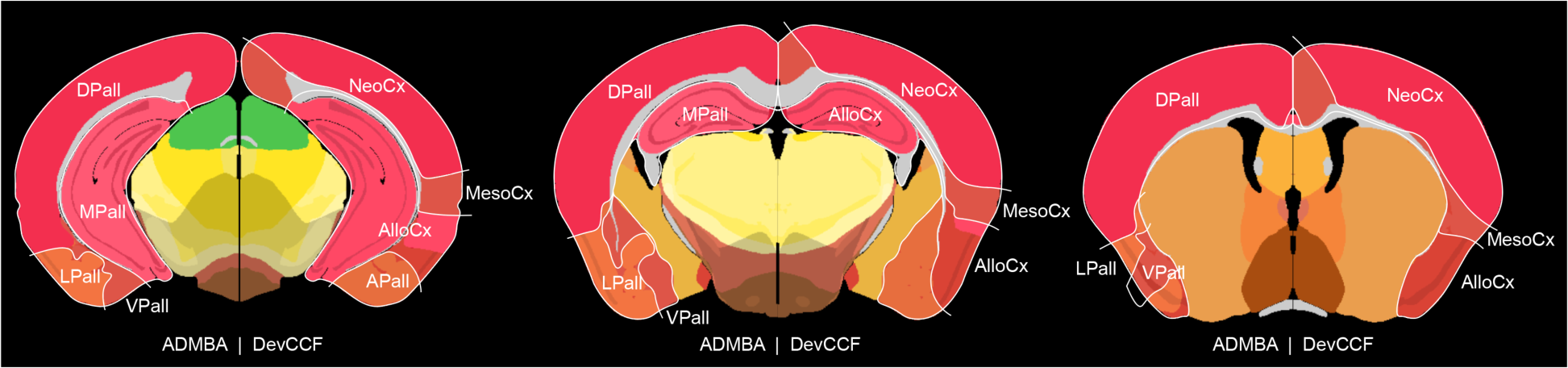
ADMBA pallium compared to concentric ring topology. Coronal slices comparing the pallium as segmented by ADMBA ontology (left hemisphere) and DevCCF ontology (right hemisphere). ADMBA ontology divides the pallium as dorsal (DPall), medial (MPall), lateral (LPall), and ventral (VPall). In contrast, concentric ring topology represented by the DevCCF divides the pallium into the neocortex (NeoCx), Mesocortex (MesoCx), allocortex (AlloCx), and Pallial Amygdala (APall).

**Extended Data Figure 2.**
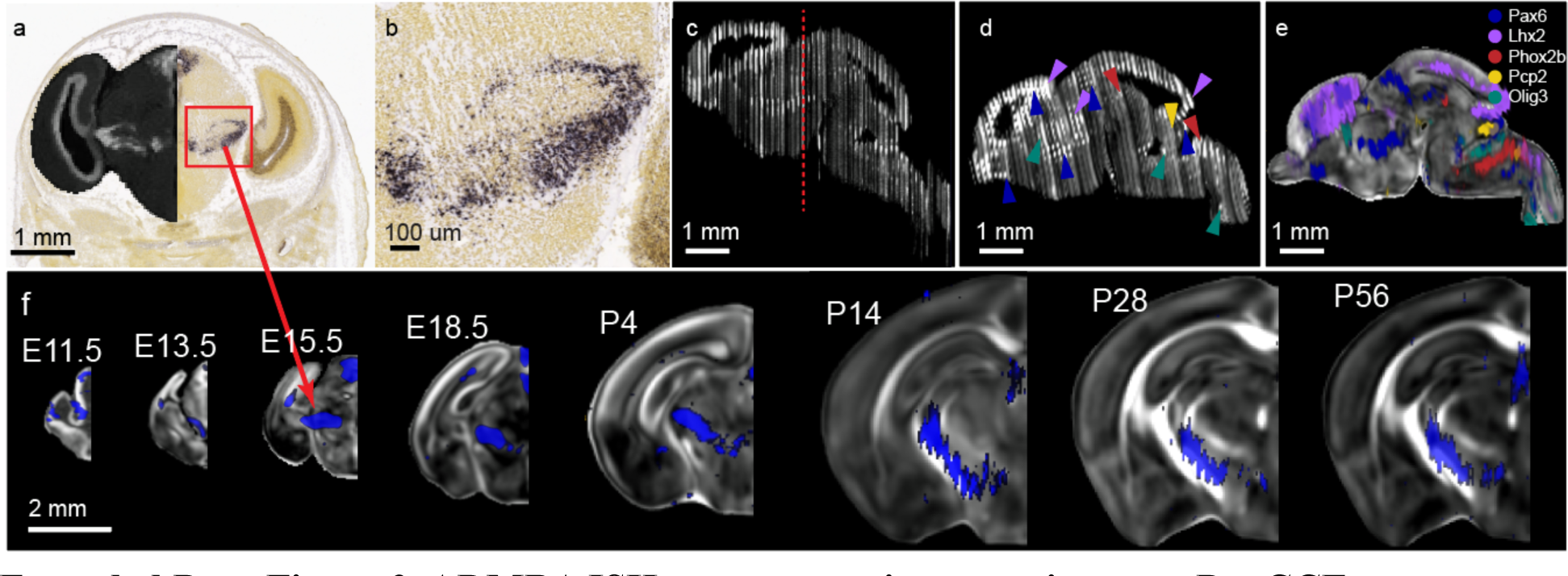
ADMBA ISH gene expression mapping onto DevCCF. (a) E15.5 ISH Pax6 slice (a) from the Allen Institute Developing Mouse Brain Atlas experiment ID 100051660. Left hemisphere overlay is dark contrast generated from inverted image intensity with greyscale color map. Red box represents location of (b). Red arrow indicates identical data registered to DevCCF E15.5 template in (e). (b) high magnification image of Pax6 expression. (c) Inverted intensity image sagittal reconstruction from coronal slices for a single subject containing ISH experiments for Pax6, Lhx2, Phox2b, Pcp2, and Olig3. Red dotted line represents (a,b) slice location. (d) Sagittal reconstruction in (c) after filling missing slices with bSpline interpolation and slice-by-slice alignment to DevCCF E15.5 template. Arrows indicate high gene expression by color. Blue: Pax6, purple: Lhx2, red: Phox2b, yellow: Pcp2, green: Olig3. (e) Localized gene expression of Pax6, Lhx2, Phox2b, Pcp2, and Olig3 (d) overlayed on DevCCF E15.5 MRI FA template. (f) Coronal view of 2D E11.5, E13.5, E15.5, E18.5, P4, P14, P28, and P56 (left to right) Pax6 ISH gene expression signal reconstruction overlayed on 3D DevCCF MRI FA templates.

**Extended Data Table 1.** Symmetric Template Sample Sex. Number of male, female, and unknown sex samples in MRI and LSFM templates at each age.

| Age | MRI Samples (Females) | LSFM Samples (Females) |
| --- | --- | --- |
| **P56** | 10 (5 Female) | 6 (3 Female) |
| **P14** | 14 (5 Female) | 10 (5 Female) |
| **P4** | 10 ($\geq 0$ Female, 9 unknown) | 8 (4 Female) |
| **E18.5** | 8 (3 Female) | 9 (4 Female) |
| **E15.5** | 9 ($\geq 2$ Female, 4 unknown) | 8 (4 Female) |
| **E13.5** | 9 ($\geq 2$ Female, 2 unknown) | 10 (5 Female) |
| **E11.5** | 6 ($\geq 1$ Female, 1 unknown) | 10 (5 Female) |

**Extended Data Table 2.**
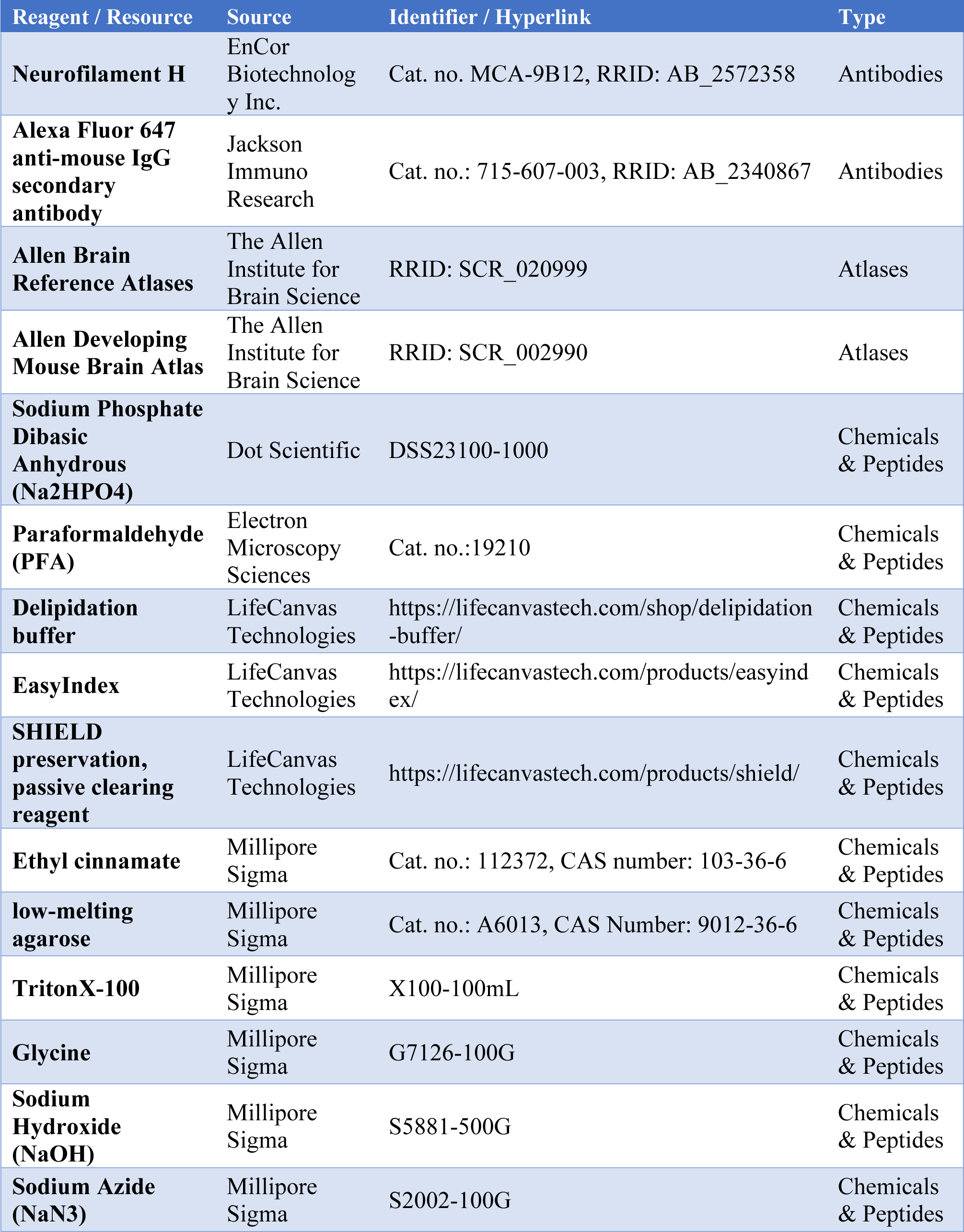
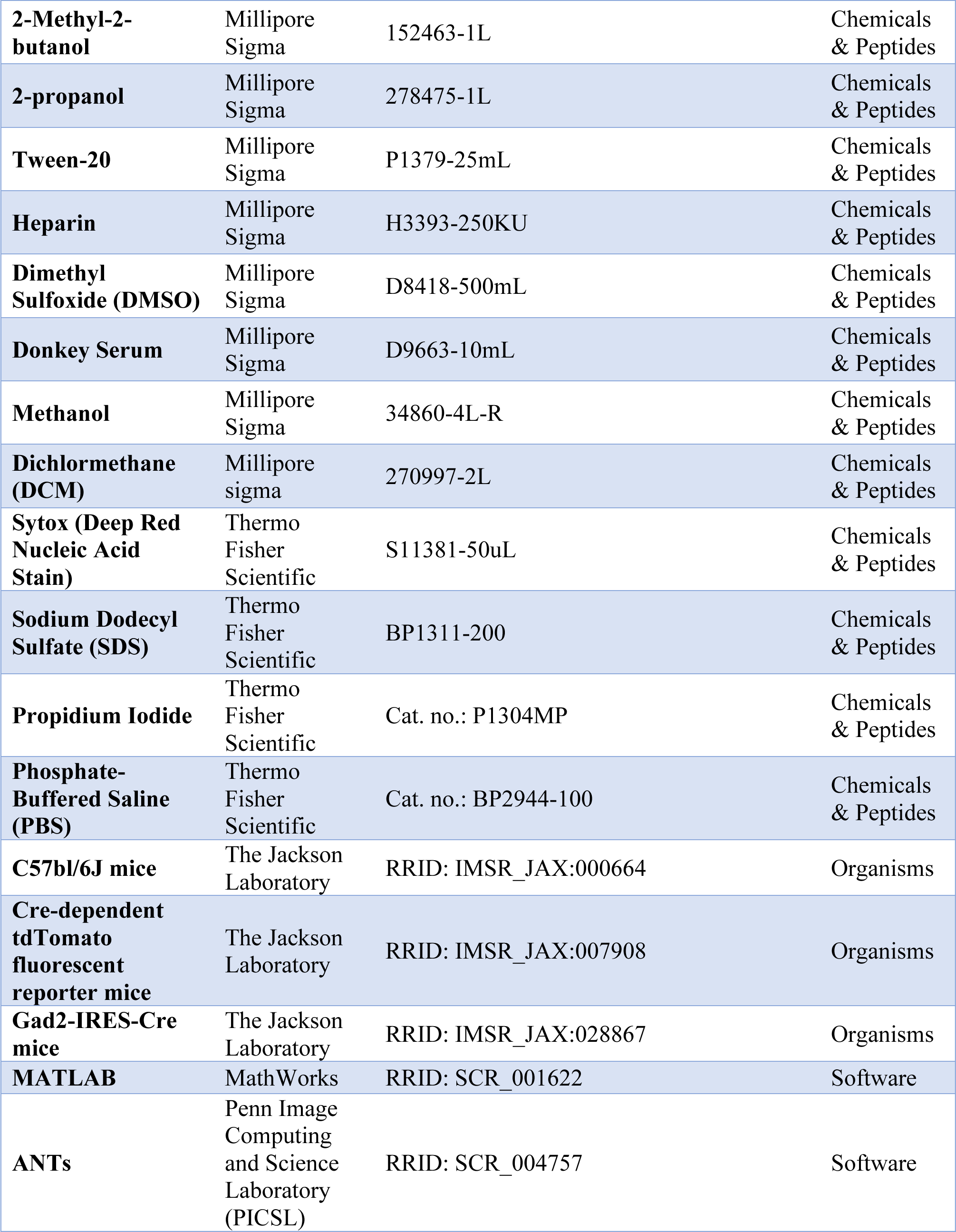
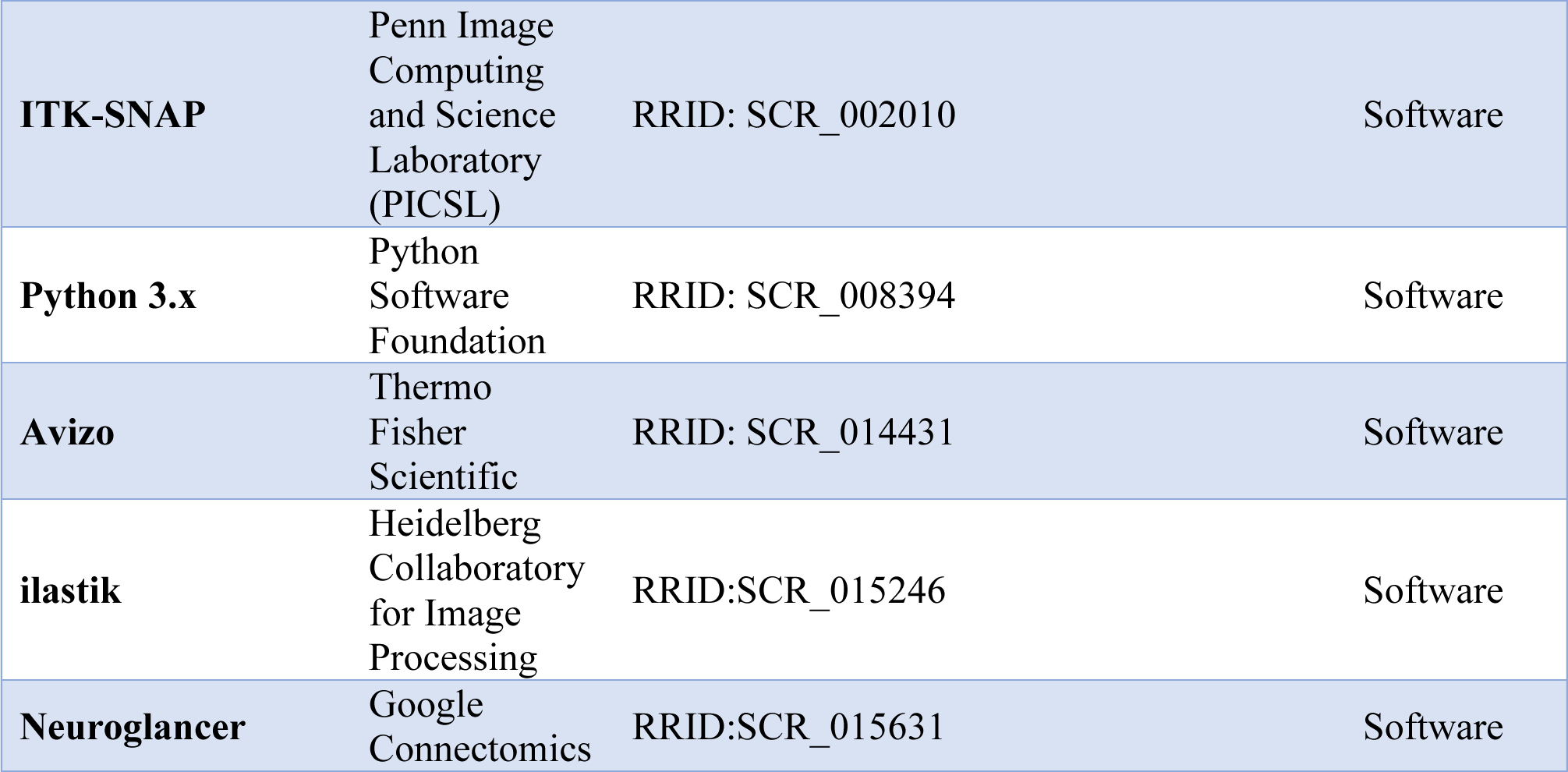
Key Resource List. List of important antibodies, atlases, chemicals, peptides, experimental animal models, and software used to develop the DevCCF.

**Supplementary Table 1. DevCCF ontology structure: See Microsoft Excel Document**

DevCCF ontology structure contains details for each region including DevCCF and ADMBA ID, name, acronym, parent, and color.

